# Updated site concordance factors minimize effects of homoplasy and taxon sampling

**DOI:** 10.1101/2022.09.26.509549

**Authors:** Yu K. Mo, Robert Lanfear, Matthew W. Hahn, Bui Quang Minh

## Abstract

**Motivation:** Site concordance factors (sCFs) have become a widely used way to summarize discordance in phylogenomic datasets. However, the original version of sCFs were calculated by sampling a quartet of tip taxa and then applying parsimony-based criteria for discordance. This approach has the potential to be strongly affected by multiple hits at a site (homoplasy), especially when substitution rates are high or taxa are not closely related.

**Results:** Here, we introduce a new method for calculating site concordance factors. The updated version uses likelihood to generate probability distributions of ancestral states at internal nodes of the phylogeny. By sampling from the states at internal nodes adjacent to a given branch, this approach substantially reduces—but does not completely abolish—the effects of homoplasy and taxon sampling.

**Availability and implementation:** Updated sCFs are implemented in IQ-TREE 2.2.2. The software is freely available at https://github.com/iqtree/iqtree2/releases.

## 1 Introduction

Site concordance factors (sCFs) are a straightforward way to measure concordance and discordance in phylogenomic data (Minh et al. 2020a). They are analogous to the standard gene concordance factors (Baum 2007), but can be calculated using all parsimony informative sites. Given a branch of interest in a phylogeny, the sCF indicates the proportion of informative (“decisive”) sites concordant with that branch. The sCF can be calculated per gene or genome-wide (i.e. from a concatenated alignment), and is especially useful when gene trees are not individually informative. Due to their ease of use and efficient implementation in IQTREE (Minh et al. 2020b), sCFs have been widely used.

Originally, the sCF was calculated by sampling quartets of tip states and asking whether the four states at the tips were concordant or discordant with a given internal branch using parsimony. However, two related factors can cause this approach to be inaccurate. First, because it is based on parsimony, homoplasy along any two branches will reduce the sCF, even when there is no discordance. Second, because quartets are sampled from across a tree, the presence of more distantly related taxa can decrease the sCF of any given branch because of increased homoplasy. Both factors will lead to artefactually lower values of sCFs.

Here, we introduce an updated version of site concordance factors that improves accuracy. To reduce the effect of homoplasy, we use likelihood to generate probability distributions of ancestral states at all internal nodes of the phylogeny. Calculations of sCFs are then made by sampling from these states at adjacent nodes in the tree. We show that the new approach is much more accurate, and is relatively unaffected by homoplasy and the presence of distantly related taxa.

## 2 Implementation and application

### 2.1 Formulation of a new sCF

Given a species tree and an alignment from the same set of taxa, we start by calculating the probability distribution of ancestral states for all sites at all internal nodes of the tree. The states at an internal node are determined by the node’s decedents only. This calculation is implemented for all nucleotide and amino acid models in IQ-TREE2. Concordance factors are calculated for each internal branch of the species tree independently. To reduce the effect of homoplasy, the updated sCF focuses on sampling states from the four nearest non-connected nodes of a specific branch, whether they are external or internal nodes. For example, to calculate the updated sCF of branch **x** in Fig. 1a, we sample a quartet of states from the probability distributions of ancestral states at nodes *a, b, c*, and *d*.

**Fig 1.**
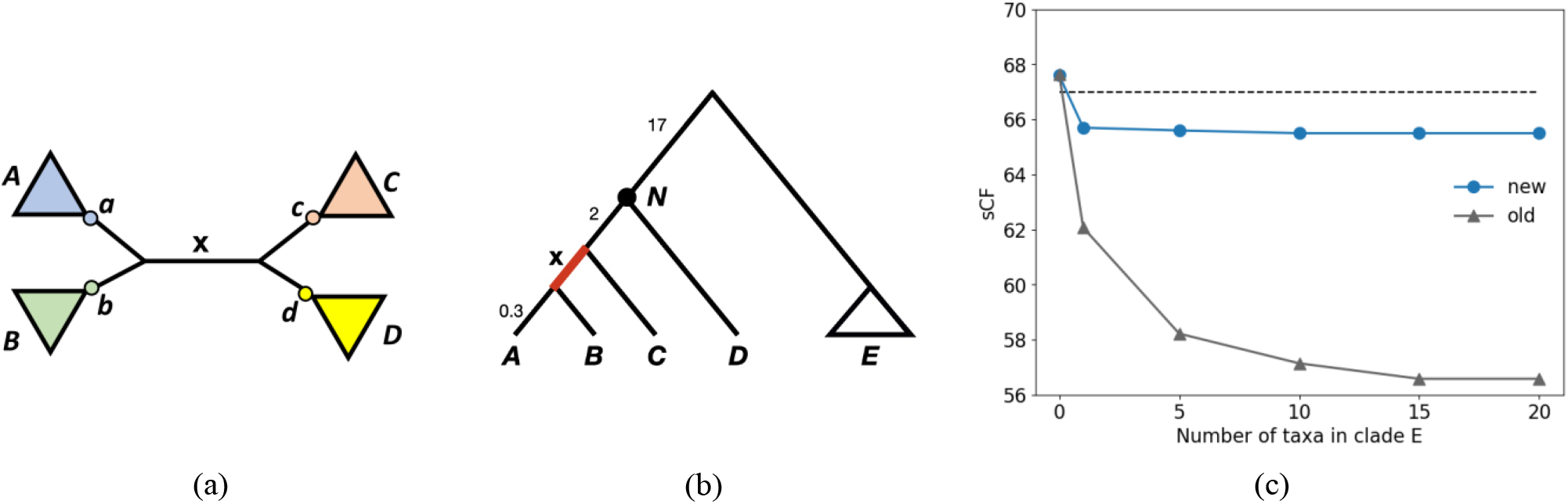
(a) A quartet will be sampled from states at internal nodes *a, b, c*, and *d* in the new sCF, but from tip taxa within clades *A, B, C*, and *D* in the old sCF. (b) The species tree used for simulation. sCFs are always calculated for branch **x** (highlighted in red), whose length is 0.7 in coalescent units. There is always one taxon in *A, B, C*, and *D*, while the number of taxa in clade *E* is allowed to vary. Note that the tree is not to scale; branch lengths shown are in coalescent units. (c) Comparison of sCFs for branch **x** in panel (b) using two methods. The new method for calculating sCFs is less sensitive to sampling distant taxa as compared to the old version. The horizontal dashed line shows the expected level of gene tree concordance.

Given a sampled state-quartet at a particular site, we say it is decisive if it is parsimony informative. For decisive sites, the state-quartet is concordant with **x** if it supports bipartition {*a, b*}|{*c, d*} (i.e. it matches the bipartition in the species tree). It is discordant if it supports either bipartition {*a, c*}|{*b, d*} or {*a, d*}|{*b, c*}. The concordance factor (CF) can then be calculated for a single sample of states in a quartet, *q*, for each of *j* sites in an alignment:

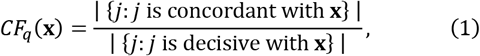

where the operator |*j*| denotes the count of sites that meet the given criterion. To account for uncertainty in ancestral states, the sCF for branch **x** is calculated by averaging over *m* sampled quartets:

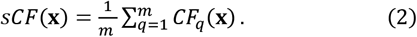

If there is one tip in a clade, the observed state at the tip is used. In the previous version of the statistic, we sampled observed states from the tips of taxa contained within clades *A, B, C*, and *D*. Therefore, the updated sCF is equivalent to the previous version for a four-taxon tree, with one tip in each clade.

### 2.2 Effect of homoplasy and taxon-sampling

In order to highlight the advantages of the new method for calculating sCFs, we simulated data with the species tree shown in Fig. 1b. Coalescent simulations were run by varying the number of taxa in clade *E*: 0, 1, 5, 10, 15, and 20. For each simulation condition, 1,000,000 trees were simulated in *ms* (Hudson 2002), and sequences were generated with *Seq-Gen* (Rambaut and Grassly 1997) setting the population mutation parameter *θ*=0.005.

The simulation is intended to demonstrate the effect on the sCF of adding more taxa to clade *E* (Fig. 1c). More taxa in clade *E* results in more quartets that sample a tip from this clade using the original sCF calculation; because of the long branch length leading to this clade, homoplasy occurs in any comparison that includes one of these tips. However, none of these additions should affect the true amount of concordance at branch **x**, as this branch length is unchanged regardless of the number of taxa in clade *E*. As more taxa are added to clade *E*, the old version of the sCF falls in value because more of the sampled quartets include a taxon from clade *E*, leading to a larger influence of homoplasy on the estimated sCF (Fig. 1c). In contrast, the new sCF does not sample tips in clade *E*: instead, it samples ancestral states from internal node *N*, which will remain strongly influenced by the observed states in taxon *D*, regardless of the number of taxa in clade *E*. As a result, it is less sensitive to the addition of distant taxa and gives a more accurate estimate of the true level of concordance. Furthermore, adding taxa to clade *E* should improve the estimate of the ancestral state at node *N* in the new approach, likely providing increased accuracy of sCFs.

Note, however, that the new version of the sCF does not solve all problems due to homoplasy. When there are no taxa in clade *E*, the two approaches are equivalent (Fig. 1c). This occurs because there are no internal nodes to sample from, and so there is no advantage of the new approach. Moreover, the addition of any taxa at all to clade *E* does cause the new sCF to go down slightly, because homoplasy still affects the distribution of states inferred at node *N*. However, additional taxa beyond the first one do not cause a further decline in the new sCF.

## 3 Summary

- We developed and implemented a new way to calculate site concordance factors based on likelihood calculations of ancestral states.
- The updated sCF statistic is much less sensitive to long branches or outlier taxa, although the problem of homoplasy is not completely removed.
- The new method is implemented and available in the latest release of IQ-TREE 2 (Minh et al. 2020b).

## Acknowledgements

We are extremely grateful to Dan Vanderpool for originally uncovering the problems addressed here.

